# A biophysically constrained brain connectivity model based on stimulation-evoked potentials

**DOI:** 10.1101/2023.11.03.565525

**Authors:** William Schmid, Isabel A. Danstrom, Maria Crespo Echevarria, Joshua Adkinson, Layth Mattar, Garrett P. Banks, Sameer A. Sheth, Andrew J. Watrous, Sarah R. Heilbronner, Kelly R. Bijanki, Alessandro Alabastri, Eleonora Bartoli

**Affiliations:** Department of Electrical and Computer Engineering, Rice University, 6100 Main Street, Houston 77005, Texas, USA; Department of Neurosurgery, Baylor College of Medicine, 1 Baylor Plaza, Houston 77030, Texas, USA

**Keywords:** brain connectivity, intracranial recordings, single-pulse electrical stimulation, pulse-evoked potentials, 3D conductivity model, tractography

## Abstract

**Background:** Single-pulse electrical stimulation (SPES) is an established technique used to map functional effective connectivity networks in treatment-refractory epilepsy patients undergoing intracranial-electroencephalography monitoring. While the connectivity path between stimulation and recording sites has been explored through the integration of structural connectivity, there are substantial gaps, such that new modeling approaches may advance our understanding of connectivity derived from SPES studies.

**New Method:** Using intracranial electrophysiology data recorded from a single patient undergoing sEEG evaluation, we employ an automated detection method to identify early response components, C1, from pulse-evoked potentials (PEPs) induced by SPES. C1 components were utilized for a novel topology optimization method, modeling 3D conductivity propagation from stimulation sites. Additionally, PEP features were compared with tractography metrics, and model results were analyzed with respect to anatomical features.

**Results:** The proposed optimization model resolved conductivity paths with low error. Specific electrode contacts displaying high error correlated with anatomical complexities. The C1 component strongly correlates with additional PEP features and displayed stable, weak correlations with tractography measures.

**Comparison with existing methods:** Existing methods for estimating conductivity propagation are imaging-based and thus rely on anatomical inferences.

**Conclusions:** These results demonstrate that informing topology optimization methods with human intracranial SPES data is a feasible method for generating 3D conductivity maps linking electrical pathways with functional neural ensembles. PEP-estimated effective connectivity is correlated with but distinguished from structural connectivity. Modeled conductivity resolves connectivity pathways in the absence of anatomical priors.

## 1. Introduction

As stimulation-based approaches for the treatment of refractory neurological and psychiatric disorders emerge as promising therapeutics, mapping human brain connectivity using electrophysiological techniques represents an integral objective in both scientific and clinical research settings (Sheth et al., 2022). The examination of brain networks and information processing spans across structural, functional, and effective connectivity domains. Recruiting the appropriate networks in an optimal manner remains an ongoing goal of stimulation-based therapies, and hinges on an advanced understanding of the intricate anatomical organization of brain areas and knowledge of the structure-function relationships. Brain networks likely represent spatially dispersed yet functionally and structurally interconnected nodes facilitating cellular communication. The clinical need for intracranial recordings creates unique research opportunities, delivering exceptional spatial and temporal resolution to provide critical insights into human brain electrophysiology and functional mapping (Keller et al., 2014). Here, we leverage intracranial electrophysiology in the construction of a three-dimensional conductivity model capable of identifying the connectivity path between stimulation and recording sites in the human brain without need for prior anatomical information.

The use of single-pulse electrical stimulation (SPES) is common clinical practice for the identification of seizure propagation patterns in medically intractable epilepsy (Boido et al., 2014). Through probing a causal influence between regions, SPES represents a method for estimating functional effective connectivity (Matsumoto et al., 2017). Application of low-frequency milliampere current pulses through electrode contacts distributes energy to the electrode-tissue interface (contact into neural tissue), disseminating its effects across associated networks and allowing estimation of electrical brain connectivity with high spatiotemporal resolution. Traditionally, cortico-cortical electrical potentials (CCEPs) are evoked and measured from the cortical surface using strips or grids of electrodes (Logothetis et al., 2010; Matsumoto et al., 2004). More recently, the use of stereo-electroencephalography (sEEG) with intracranial depth electrodes has enabled the recording of neural activity across cortical, subcortical, and white matter regions. CCEP produces relatively clear response profiles across sampled brain regions due to the consistent orientation of neuronal components (Crocker et al., 2021). However, surgical insertion of sEEG electrode leads at varied orientations results in differing stimulation dipoles relative to cortical axes (Prime et al., 2018). Having neurons at different orientations imparts a spectrum of response morphologies during SPES in sEEG, wherein orientations favoring the cortical axis exhibit stronger responses (Paulk et al., 2022). Recognizing this key difference, we refer to the event-related responses (ERP) to SPES recorded through sEEG as pulse-evoked potentials (PEPs).

SPES has aided in the mapping of functional brain networks, including the language network (Matsumoto et al., 2004), limbic network (Enatsu et al., 2015), and frontal lobe networks (Greenlee et al., 2007). Several ERP features are reported in SPES. A sharp early negative potential (N1 peak) occurring 10-50 ms post-stimulation, and a second slow-wave potential (N2 peak) typically occurring 50-300 ms are commonly used in CCEPs to characterize response morphology (Matsumoto et al., 2004). These ERP components may reflect anatomical or functional processing in neuronal ensembles (Kundu et al., 2020; Luck, 2014). Similar response components exist in sEEG PEPs, although they occur with higher variability, likely due to inconsistencies in neuronal morphology and dipole orientation across sampled spatial locations. To acknowledge key differences between CCEP and sEEG PEPs, we exploit the relative temporal constraints presented in N1 and N2 detection to employ a consistent and automated detection method for the identification of the first two response components in sEEG SPES, hereby labeled C1 and C2. By adopting this terminology, we preserve the value of each response element as explanatory profile metrics, while remaining agnostic to their precise correspondence to N1 and N2 CCEP components.

Comparisons of SPES with other forms of connectivity have laid a foundation for relating PEPs to established structural connectivity measures (Babaeeghazvini et al., 2021). Diffusion magnetic resonance imaging (dMRI) is a non-invasive technique that uses information on the displacement of water molecules in neural tissues to reconstruct the white matter pathways mediating anatomical connections in the brain (Le Bihan and Iima, 2015). Structural connectivity can be quantified using probabilistic tractography algorithms to provide an estimate for the strength of the association between nodes. In tractography, the number of streamlines is considered a proxy for axonal projections and is commonly reported as the primary outcome measure. Tractography-based connectomes are increasingly used for image-guided surgical planning (Essayed et al., 2017; Noecker et al., 2018). However, the inability of tractography algorithms to accurately recapitulate all elements of underlying anatomical connectivity means that additional measures of connectivity are sorely needed (Grier et al., 2020; Jbabdi and Johansen-Berg, 2011; Thomas et al., 2014). Still, recent literature highlights cross-modality similarities with PEPs across several brain regions (Adkinson et al., 2022; Crocker et al., 2021). Despite reporting distinct types of connectivity, the correlation between tractography and SPES-induced PEPs suggests that the two modalities may share the ability to report information on pathways supporting network connectivity.

As a conductive medium, the human brain facilitates the transmission of information in part via electrical signaling. Within biological tissue, conductivity constitutes the movement and density of ionic content within an electric field, and such values are tissue-type dependent (McCann et al., 2019; Miklavčič et al., 2006). Estimates of conductivity in neural tissue are used widely in transcranial magnetic stimulation (TMS), as the stimulation-induced electric field is influenced by fluctuations in conductivity at interfaces between gray matter, white matter, cerebrospinal fluid (CSF), and the skull (Niu et al., 2023). In intracranial electrophysiology, conductivity is one of the several important biophysical properties considered when calculating the electric field created by stimulation (Alonso et al., 2023; Carvallo et al., 2019; Howell et al., 2019). In the present paper, we introduce a novel method for estimating three-dimensional *conductivity* maps based solely on intracranial recordings during SPES. Our proposed model is naïve to anatomical priors, removing the need for access to dMRI, data, which remains limited to institutions with sufficient technical expertise. We employ a passive propagation of electric potentials from the stimulated electrodes to the recording electrodes and resolve a 3D conductivity solution that fits the potential values measured by the C1 PEP peaks. In this way, we identify and reconstruct conductivity pathways connecting nodes of functional networks. Importantly, this model allows us to identify electrical pathways that are consistently engaged in order to directly compare PEPs with tractography estimates.

## 2. Materials and Methods

The following section describes the electrophysiological recordings, stimulation, and the features extracted to capture the pulse-evoked potential (PEP) responses across the recording locations and the diffusion magnetic resonance imaging (dMRI) protocol.

### 2.1 Electrophysiological Data

#### 2.1.1 Human subject

One human subject (40y old female) provided informed consent to participate in this study during ongoing intracranial epilepsy monitoring with stereo-electroencephalography (sEEG) at our institution. Experimental procedures were conducted in accordance with the policies and principles outlined in the Declaration of Helsinki and were approved by the Institutional Review Board at Baylor College of Medicine (H-18112). The patient had no prior surgery, and brain imaging was unremarkable. Single pulse electrical stimulation (SPES) experiments were conducted one day following electrode implantation the patient was still on anti-epileptic medications and interictal epileptic activity was at its minimum.

#### 2.1.2 sEEG probes

The subject underwent invasive surgical implantation of 14 depth probes, with a total number of 152 electrode contacts spanning various anatomical locations across right frontal and bilateral temporal regions (Figure 1a, b). Electrode contacts ranged from 8-16 per probe, with an median of 10 electrode contacts per probe. The sEEG probes had either a 0.8 mm diameter with 8-16 electrode contacts and a 3.5 mm center-to-center distance (PMT Corporation, MN, USA) or a 1.28 mm diameter with 9 recording contacts and a 5.0 mm center-to-center distance between contacts (AdTech Medical Instrument Corporation, WI, USA).

**Figure 1.**
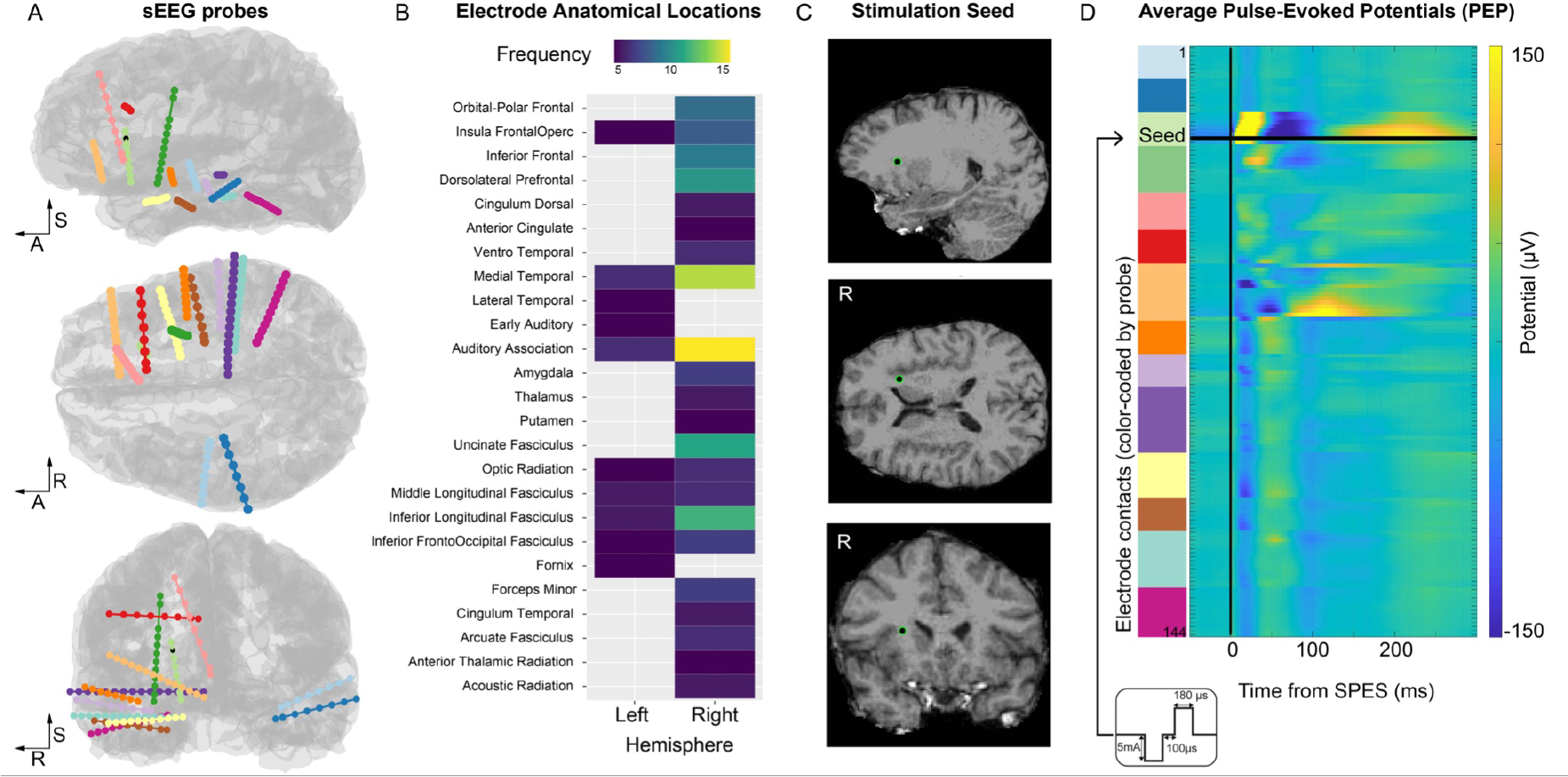
Overview of anatomical locations and stimulation protocol. **A**. All electrode contacts (144 electrodes represented as spheres) along each sEEG probe (14 probes, color-coded) are displayed on the sagittal, axial, and coronal views of the cortical surface. The R-A-S axis (right, anterior, superior) are included for reference. Electrode shape and size are distorted and exaggerated for visualization purposes. The electrode used as the stimulation seed in Experiment 1 is represented by a black sphere with a green outline. **B**. Overview of the electrode anatomical locations color-coded by the number of electrodes in each anatomical parcellation (based on the HCPex modified atlas). **C**. Detailed view of the location of the stimulation seed in Experiment 1: the electrode is located in the white matter adjacent to the anterior insular cortex. **D**. Average pulse-evoked potential (PEP) across all electrodes in response to single-pulse electrical stimulation (SPES) delivered to the stimulation seed. The inset shows the schematic of the SPES waveform and parameters.

#### 2.1.3 Anatomical localization of electrodes

Locations of electrode contacts were determined by employing the intracranial Electrode Visualization software pipeline, iELVis (Groppe et al., 2017). Briefly, the acquired post-operative clinical CT image was co-registered to the pre-operative T1 anatomical MRI scan using the linear image registration tool (FLIRT) as part of the Functional Magnetic Resonance Imaging for the Brain Software Library (FMRIB) (Jenkinson and Smith, 2001). Next, the location of each electrode contact was manually identified on the CT-MRI overlay in BioImage Suite (Papademetris et al., 2006). The *xyz* coordinates for each electrode represent their position in space (mm) in an R-A-S convention (increasing values of x point to the Right, y to Anterior, and z to Superior, with an origin of the central voxel of the T1 volume). Coordinate locations of electrode contacts were extracted for anatomical labeling purposes. Contacts residing in cortical and subcortical regions were assigned anatomical locations according to a modified version of the volumetric HCPex atlas (Huang et al., 2022), while the XTRACT HCP probabilistic tract atlas (Warrington et al., 2020) was used for labeling contacts situated in white matter (Figure 1b). In the case that a contact existed outside of the parcellations, such as superficial white matter, the most likely parcellation estimate was detected based on spatial proximity.

Additional anatomical information was obtained by quantifying electrode properties based on the proximity to the cortical surface and white-gray boundary surface, each reconstructed from the T1 using FreeSurfer (version 6.0; (Dale et al., 1999; Fischl, 2012)). First, for each electrode location, we computed its minimum distance to the pial surface and to the white-gray boundary; we obtained the cortical parcellation estimate for that closest cortical surface point based on the Destrieux Cortical Atlas (Destrieux et al., 2010). The atlas contains 76 cortical parcellation labels and each cortical surface point is assigned a label by mapping the parcellation estimates on the individual cortical surface. To quantify the complexity of nearby anatomy while constraining the maximum search distance allowed, we detected all cortical parcellation estimates within a 5 mm search radius around each electrode and ranked the labels by their proportion (e.g. one electrode is labeled as 91% superior temporal gyrus and 9% superior temporal sulcus). Lastly, from the ranked labels, we computed the likelihood of each electrode being located near a sulcus, near a gyrus, or a mixture of both by reducing the labels in these 3 categories (e.g. the example above would be 91% gyrus and 9% sulcus).

#### 2.1.4 Electrophysiological stimulation

We employed a monopolar cathodic SPES paradigm. Biphasic pulses with 5 mA amplitude, 180 μs pulse width, and a 100 μs interphase gap were delivered to stimulated electrode contacts (stimulation seeds) using a Blackrock CereStim R96 stimulator (Blackrock Microsystems, Utah, USA). For each stimulation seed, a total of 315 single pulses were delivered with a variable inter-stimulation period ranging uniformly between 400 ms and 1.2 s. We performed a total of 3 stimulation experiments using the parameter configuration described above and sampled several anatomical locations in the right hemisphere of the subject. Stimulated regions included the anterior insula (Experiment 1, Figure 1c), the posterior thalamus (Experiment 2) and subgenual cingulate cortex (Experiment 3). Appendix A contains the anatomical visualization for the three stimulation seeds and additional anatomical information for the sampled cortical locations. The selection of stimulation seeds was motivated by both anatomical and clinical considerations (electrode contacts must not be in the suspected seizure onset zone and must be at least 10 mm from the reference electrode).

#### 2.1.5 Electrophysiological recording and preprocessing

Neural signals from the sEEG probes were recorded using a 256-channel Blackrock Cerebus system (Blackrock Microsystems, UT, USA) at 30 kHz sampling rate, with a 4^th^ order Butterworth high pass filter (0.3 Hz). Signals were recorded during each stimulation experiment from all electrode contacts except for the contact being stimulated. sEEG recordings were referenced to a pre-selected electrode contact visually determined to be in white matter during the implantation procedure. An in-house, custom MATLAB script was used to process acquired stimulation data. Initially, the quality of sEEG signals were visually inspected for line noise, excessive recording artifact, jaw contraction contamination, and interictal epileptic spiking. Electrode contacts displaying poor signal quality were excluded from analysis. Following the application of exclusion criteria, a total of 144 electrodes (out of 152) were used in signal-based analyses. The neural signal around each SPES onset was identified, and a window of interest was defined between 300 ms pre-stimulation and 300 ms post-stimulation. An average waveform (one for each electrode contact) was calculated across the 315 individual SPES response profiles (Figure 1d). This average waveform was extracted and used in the identification of specific PEP features. The same procedure was repeated for the different stimulation experiments.

#### 2.1.6 Extracting PEP (pulse-evoked potential) features

Several features of PEPs were calculated in the present experiment. Below, we describe features obtained from the automatic detection of the C1 and C2 waveform components (Peak Amplitude, Latency, Area, Peak-to-peak Distance) and features calculated from the overall average waveform (i.e., not requiring detection of the components, Maximum Amplitude and Root Mean Square of Average Shape).

##### 2.1.6.1 Automatic Detection of C1-C2 component characteristics

The average waveform recorded at each channel location was high-pass filtered (cutoff frequency of 300 Hz), and the filtered signal was subtracted from the average signal to obtain a smoothed average waveform with no high-frequency artifacts that could affect the automatic peak detection. The maximum and minimum values within the temporal window 10-80 ms after SPES were identified, corresponding to the peak of the C1-C2 components. This approach detects C1 as the first component (regardless of polarity), and C2 as the second component (opposite polarity with respect to the C1). The inflection points preceding and following the identified peaks were used to determine the initiation and termination of each component. Then, the linear segment connecting the inflection points was calculated and used to delimit peak boundaries (Figure 2).

**Figure 2.**
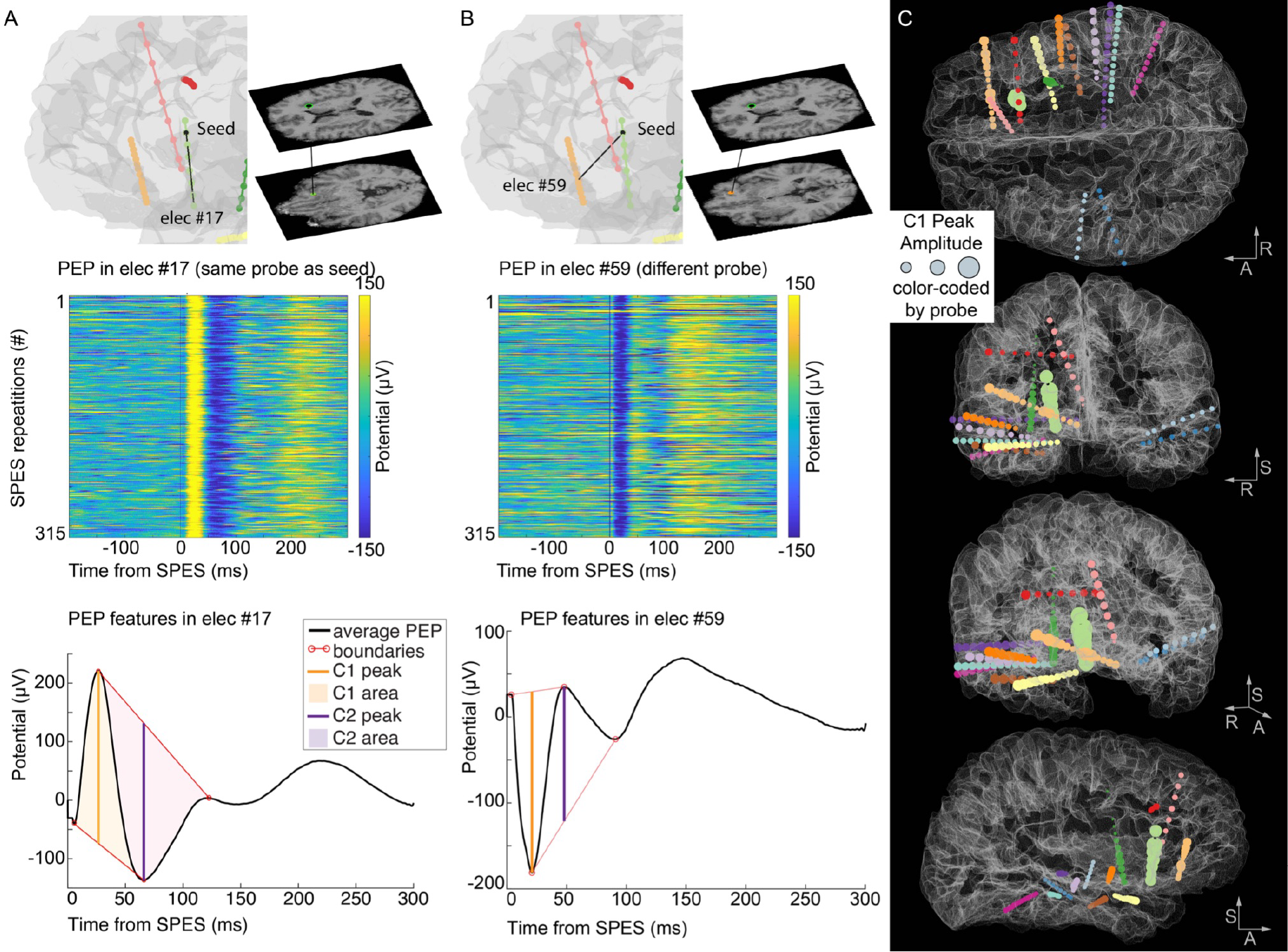
Example of PEP features and their distribution across electrode locations. **A**. On the top, a visualization of the relative position between the stimulation seed for Experiment 1 and an electrode on the same probe (elec #17). The middle panel displays the PEPs recorded from that electrode in response to each SPES delivered at the stimulation seed, showcasing the consistency of evoked potentials recorded from electrode #17. Below, the PEP features (C1 and C2 peaks, their amplitude, their area, and the peak boundaries) are represented on the average waveform. **B**. Same information as in A, recorded from a different electrode (elec #59). Note the opposite polarity of the PEP waveforms between the two recorded locations. **C**. The C1 Peak Amplitude is represented at each electrode location by inflating the size of the sphere by the C1 amplitude (i.e., larger spheres represent larger C1 amplitudes). The electrode locations are color-coded by the probe as in Figure 1. The stimulation seed electrode is marked by a black sphere (visible on the coronal and sagittal views). The R-A-S axis (right, anterior, superior) are included for reference.

###### C1 and C2 Peak Amplitude

Each component amplitude was quantified as the difference between the potential value at its peak and the voltage value at the linear segment during the corresponding time point. Note that the component features were calculated independent of their polarity, such that a C1 component could present as either the first positive or the first negative deflection following SPES.

###### C1 and C2 Latency

The time point at which the C1 and C2 peaks were detected with respect to the SPES onset.

###### C1 and C2 Area

The area of each component was calculated as the cumulative sum of all elements enclosed beneath the curve and the linear segment boundary.

###### Peak-to-Peak Distance

We defined peak-to-peak distance as the distance between the detected maximum and minimum peaks (C1-C2 difference).

##### 2.1.6.2 Calculation of PEP features independent of PEP components

###### Max Amplitude

The maximum potential value occurring in the 10-200 ms window following the stimulation pulse was obtained (regardless of polarity), as a proxy of the strongest collective response of a contact during stimulation.

###### RMS of Average Shape

The root mean square (RMS) technique has been used previously to characterize CCEP responses (Dionisio et al., 2019; Prime et al., 2018). Here, we evaluate the RMS of the phase-consistency calculated across SPES repetitions (RMS-Shape) in sEEG in an effort to define consistency in the morphology of response profiles. Contacts displaying high values for RMS-Shape exhibit a consistent phase across individual trials in the prescribed analysis window.

### 2.2 Diffusion Magnetic Resonance Imaging (dMRI)

#### 2.2.1 Imaging acquisition

Prior to surgical implantation of sEEG electrodes, the patient participated in anatomical imaging at the Core for Advanced Magnetic Resonance Imaging (CAMRI) at Baylor College of Medicine (Houston, TX, USA). Diffusion-weighted imaging (DWI) data was acquired on a Siemens Prisma 3T system for 2 b-values (b = 1000, 2000 s/mm^2^) with two phase-encoding directions (anterior-to-posterior and posterior-to-anterior) using a dual-echo echo-planar protocol (Repetition time (TR) = 3400 ms, Echo time (TE) = 85.8 ms, Voxel size: 1.5 mm isotropic, Multiband acceleration factor = 4, 99 slices). A total of 7 interleaved non-diffusion-weighted (b0) volumes were collected and 92 diffusion-sensitizing gradient directions were applied per acquisition.

#### 2.1.2 DWI processing and diffusion model fitting

DWI data was processed using FSL’s diffusion toolbox (Jbabdi et al., 2012b). Initially, image pairs acquired with reverse polarities (distortions in opposite directions) were aligned across the time domain. From the merged DWI data, all 14 b0 volumes were used to estimate the susceptibility-induced field and correct for resultant geometric distortion (Andersson et al., 2003). Additionally, distortions introduced by subject movement and eddy-current induced distortions caused by rapid diffusion gradient application were corrected using *eddy* software (Andersson and Sotiropoulos, 2016). A multi-shell diffusion model based on Markov Chain Monte Carlo sampling was then fitted to the preprocessed 4-dimensional DWI data to estimate intra-voxel fiber orientations (Jbabdi et al., 2012a). This approach uses anisotropic tensors in isotropic backgrounds to model free diffusion and axonal tracts, respectively.

#### 2.1.3 Probabilistic tractography

To reconstruct white matter pathways and estimate structural connectivity non-invasively, probabilistic tractography was employed. To generate regions of interest (ROIs) for electrode contacts, coordinate locations of each contact were extracted to create 2 mm radius spherical masks. Spherical masks were moved from native pre-operative structural to native diffusion space via linear transformation. Generated ROIs are representative of the stimulation field of electrode contacts. Masks from stimulation seed locations were used as seeds and tractography was initiated and run iteratively from every stimulation seed to each electrode contact target mask. Tractography was conducted using the default parameters of FSL’s *probtrackx* algorithm with a distance correction feature applied (samples = 5000, curvature threshold = 0.2, step length = 0.5 mm, subsidiary fiber volume fraction threshold = 0.01, loopcheck termination) (Behrens et al., 2007, 2003). A cerebrospinal fluid (CSF) exclusion mask was implemented to prevent erroneous tracking in CSF, and each target electrode was considered both a waypoint and termination mask to enable estimation of the connectivity between seed and target regions (Zhang et al., 2001).

##### #Streamlines

The number of streamlines, representing connectivity likelihood, was then extracted for each seed-target pair from the *waytotal* output and used in statistical evaluation against derived PEP features. To reduce skewness, a logarithmic base 10 transformation was applied to the raw streamline data count.

##### Streamline Length

For every streamline found between the each seed-target pair we extracted its path length as a measure of distance (in mm) between voxel locations of the seed and target region. We averaged the path lengths obtained across all the streamlines obtained for each seed-target pair.

## 3. Calculations

### 3.1 Conductivity Model

We use COMSOL Multiphysics® version 6.1 (Stockholm, Sweden), a Finite Element Method (FEM) software, to build and solve a model of brain connectivity based on the C1 SPES data. Rather than using statistical and machine learning methods to estimate functional connectivity from SPES data or other neurophysiological biomarkers such as EEG (Rossini et al., 2019), methods which may struggle in terms producing readily-interpretable descriptions of connectivity for a clinical audience (Murdoch et al., 2019), our FEM model solves a current conservation problem

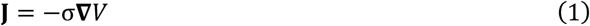

for the electrical potential *V* to construct a visual analog of connectivity in the form of an electrical conductivity map σ. Here, **J** is the current density.

Our approach provides a physically consistent conductivity map in accordance with the experimental data, while describing a passive system (i.e., it does not account for voltage increases along electric current paths). As a result, while the absolute values of the calculated conductivities are not representative of local electrical phenomena, the relative values across the probed brain locations determine functional connectivity compatible with neural signal pathways.

We estimate the spatial conductivity map σ through topology optimization of a domain of electrically conductive material. Topology optimization methods seek to generate the optimal structural layout for a system based on some design objective. Originally proposed as the continuous generalization of the discrete problem of shape optimization, topology optimization finds the best distribution of material within a given geometry using a continuous density model control variable (Bendsøe, 1989). These techniques have long been applied to optimize multiscale systems from a structural mechanics perspective in diverse fields such as mechanical engineering, aerospace engineering, architecture, and medical technology (Wu et al., 2021). More recently, topology optimization has been applied to electromagnetic metasurfaces as well (Zhao et al., 2022). Here, we present a novel application of topology optimization in neuroscience, where we produce simple physical models which aid in visualizing and understanding functional connectivity in the brain.

Before considering full 3D models to directly compare to the C1 Peak Amplitude values, we built simplified 2D models to investigate numerical procedures and parameters estimation with reduced computational expense and to provide clear descriptions of the modeling approach (Figure 3a shows a diagram of our FEM model in 2D). Given spatial coordinates of the sEEG electrodes, we construct a simple geometry containing circular (2D) or spherical (3D) electrodes of radius 0.65 mm within a circular or spherical domain of radius 100 mm. In both cases, we place an electrical ground (*V* = 0 µV) on the exterior boundary of an extended domain (representing the probed brain regions) and a constant electrical potential *V*_0_ on the boundary of a single electrode domain deemed the source electrode (stimulation seed). The electrodes are modeled as made of metallic material with large electrical conductivity 6 × 10^7^ S/m. The electrical conductivity in the domain, σ = σ(*x, y*), is a function of position determined iteratively by topology optimization of a density model. For the optimization the density model control variable is an artificial volume factor, 0 ≤ θ_c_ ≤ 1, defined across the domain and varied during the optimization process described below.

**Figure 3.**
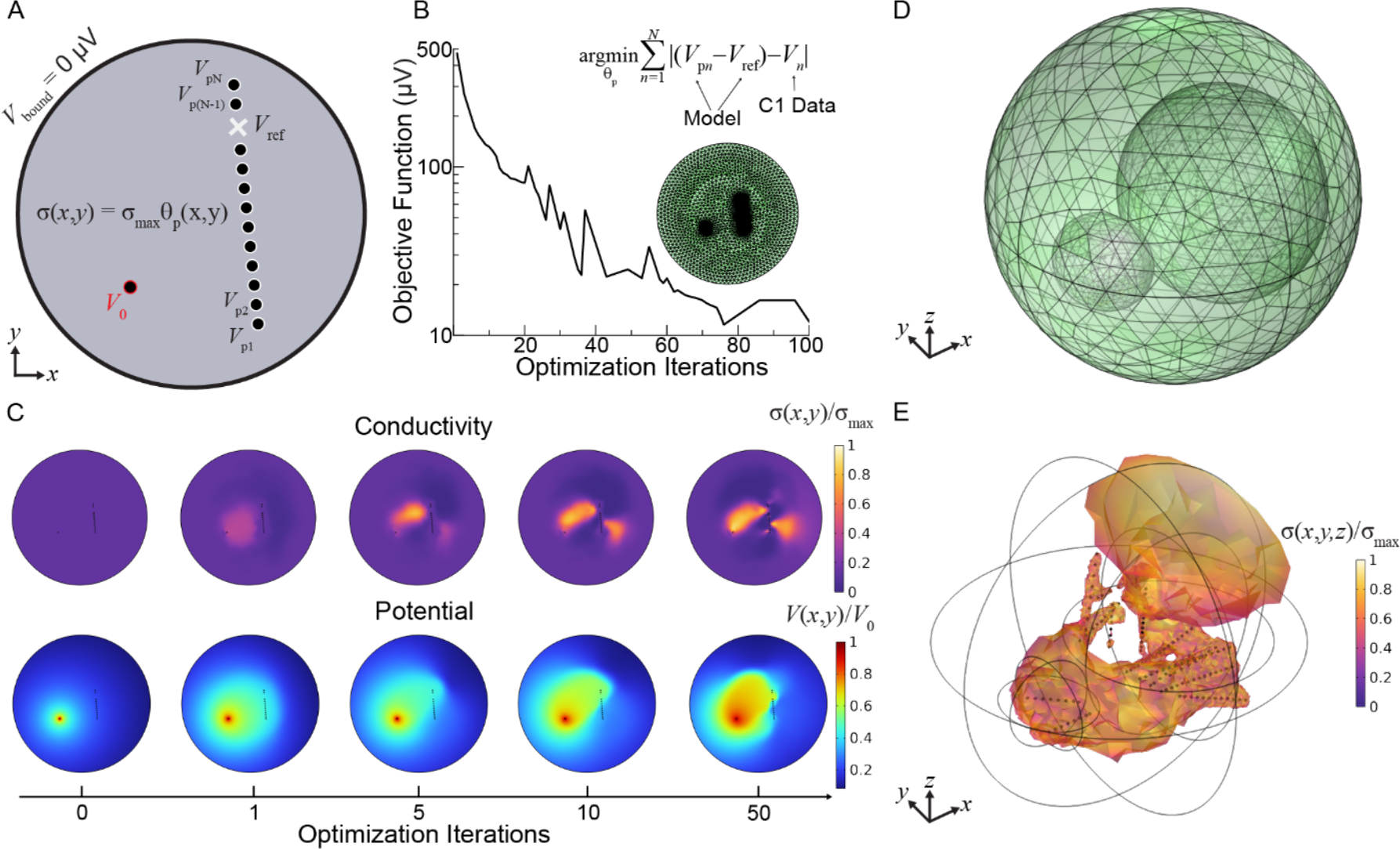
Diagram of 2D Finite Element Method (FEM) model of brain connectivity. **A**. Geometry consists of a circular domain (radius 100 mm) of material with electrical conductivity given by the penalized density model θ_p_ from topology optimization scaled by a maximum conductivity σ_max_, and metallic electrodes (radius 0.65 mm) with electrical conductivity 6×10^7^ S/m. There are no electrical boundary conditions beyond the source potential *V*_0_ on the boundaries of the stimulated electrode and electrical ground on the boundary of the domain. The electric potential is probed at each electrode (*V*_p*n*_) in addition to the reference potential (*V*_ref_), locations dictated by clinical probe locations, to compute the objective function at each optimization solver iteration. **B**. The objective function–the sum of differences between the referenced electrode potential measurements from the model and the C1 data (*V*_*n*_)–is iteratively minimized to find the density model θ_p_ that scales the conductivity map across an FEM mesh (green). **C**. Selected 2D model solution plots over increasing optimization iterations demonstrating how the conductivity map (top) and thus the electric potential (bottom) evolves as the objective is minimized. **D**. The modeling procedure is directly extended to 3D by using spherical domains. The additional ellipsoids are part of the topology optimization domain and are only to control the growth rate of the tetrahedral mesh. **E**. Example 3D conductivity map using C1 data from 143 clinical electrodes in the objective, filtered so that only mesh locations exceeding 40% of the maximum conductivity appear for visual clarity.

To improve the computational procedure, instead of directly scaling the domain conductivity, we enforce a minimum feature size, *R*_min_, by computing a filtered control variable θ_f_ as a solution to a Helmholtz equation:

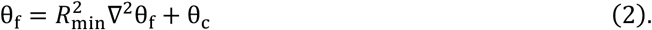

This filtering limits the complexity of the control variable map and enforces a certain degree of smoothness by smearing rough, small details into a grayscale blur. In all our models, *R*_min_ is the minimum mesh element size. This smearing effect is reduced by a hyperbolic tangent projection of θ_f_

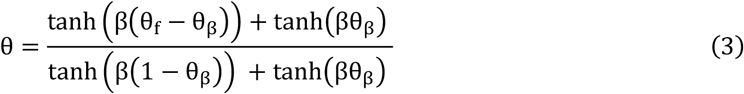

where β and θ_β_ are parameters controlling the strength of the projection. The hyperbolic tangent projection sharpens the solution but also increases the computational expense of the optimization, with larger values of β increasing both effects. Finally, inspired by material properties optimization in mechanical systems, we filter the projected control variable θ once more via the Solid Isotropic Material with Penalization (SIMP) method (Bendsøe, 1989):

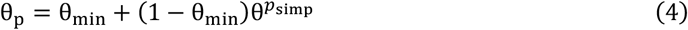

this encourages well-defined structure in the final control variable map *θ*_*p*_ by forcing the density model toward small values (voids) or large values (solid material). The minimum value, θ_min_, ensures the solution does not vanish anywhere and is set to 0.001 for all of our models. The exponent *p*_simp_ controls the strength of the penalization.

We thus define the electrical conductivity σ of the domain encapsulating the metallic electrodes as a scaling of the penalized density model

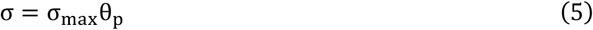

where σ_max_ is the maximum electrical conductivity. We initialize the conductivity with θ_p_ = 0.2. With the electrical conductivity now defined everywhere in our model geometry, we begin the topology optimization by solving for the electric potential *V* with Eq. 1. After the FEM solution for the potential converges, we probe the potential at each electrode: given *N* electrodes we collect electrode domain probe voltages **V**_**p**_ = [*V*_p1_, *V*_p2_, …, *V*_p*N*_]. Additionally, we probe a reference potential *V*_ref_ at a specific coordinate in space consistent with experimental data. In models based on C1 data, this reference potential is taken at the same reference location as in the clinical measurement. Then, comparing with the clinical C1 potential data, **V** = [*V*_1_, *V*_2_, …, *V*_*N*_], we evaluate the objective function:

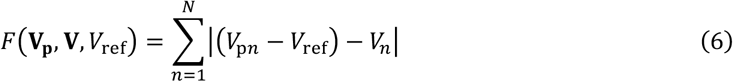

which is the sum of differences between the referenced model electrode potentials and the C1 data. With the Sparse Nonlinear OPTimizer (SNOPT) solver (Gill et al., 2005), we iteratively vary the map of θ_p_ and re-solve Eq. 1 to minimize Eq. 6. Figure 3b plots the decrease of the objective function over successive solver iterations. The final conductivity map is thus 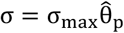 where:

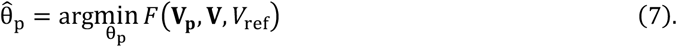

Figure 3c gives a visual illustration of the optimization process in 2D, showing how the iteratively varied conductivity map changes the solution for the electric potential at increasing iterations. In 3D, the modeling procedure is exactly the same as described above beyond the addition of the *z* spatial coordinate and the use of spherical domains. Figure 3d displays the 3D FEM mesh for the *N* = 143 electrode model used to compare with the clinical C1 data. The 3D model produces conductivity maps like the one displayed in Figure 3e.

The key parameters we varied in search of the most accurate model of the C1 data were θ_β_, β, *p*_simp_, and σ_max_. Trends in model accuracy were non-monotonic with all solver parameters. Thus, we performed a grid search over reasonable parameter values for the hyperbolic tangent projection and SIMP penalization to achieve the lowest average electrode error for a given dataset. The lowest error was achieved with moderately strong projection given by θ_β_ = 0.1, β = 8, and cubic SIMP penalization given by *p*_simp_ = 3. As we are using electrical conductivity only as a proxy for neural connectivity, the absolute value of the conductivity in S/m is not physically associated with the exchange of information between brain regions. However, on certain C1 Peak Amplitude values we did find that increasing the maximum conductivity within the range physiological neural tissue conductivities (McCann et al., 2019) —from 0.1-0.33 S/m to 1 S/m—improved average electrode error (defined as % absolute difference between C1 values and output electrode voltages) slightly without qualitatively changing the conductivity map, supporting the rationale that, despite describing a passive system, the calculated spatial variation of the conductivity values is indicative of neural activity. Regardless, tuning the parameters listed above was important in achieving the most accurate result for each dataset. It should be noted that the optimization problem as casted here is typically under constrained given the limited spatial sampling regions provided by electrode number and locations. The use of a larger number of probes in many regions can mitigate the optimization and lead to more accurate visualization of interconnected neural areas.

## 4. Results

Initially, we compare and validate different PEP features extracted from our data. We evaluate the agreement between different metrics, including those obtained with an automatic detection method to identify C1 and C2 neural components from pulse-evoked potential (PEP) responses to single pulse electrical stimulation (SPES). We then probe relationships between tractography-estimated structural connectivity and PEP metrics in order to measure effective connectivity. We select the C1 component to guide our proposed optimization model in resolving a conductivity map between the stimulation seed and electrode contact targets. The C1 Peak Amplitude values are used to construct the objective function in topology optimization and the locations of electrode contacts are utilized by the physical model to represent the probe insertions. While the electric current equations do not directly employ C1 values, the optimization solver operates iteratively to shift the solution toward C1 data, making repetitive comparisons of model values with C1 values. We lastly evaluate the model results by comparing the conductivity map with anatomical information and by identifying anatomical features that induce large voltage gradients and lead to large model errors at those locations.

### 4.1. Heterogeneity of C1 and C2 components across electrode locations

After visually inspecting and removing channels with significant biological and stimulation artifacts, the smoothed average waveform at each electrode contact was evaluated for identification of C1 and C2 response components. We found that the average waveforms are dissimilar across different electrode contacts (Figure 2a, b), particularly when considering polarity, but also overall waveform morphology. Given the variable orientation of neural structures and resultant heterogeneous polarities at electrode contacts in sEEG, we detected the C1 and C2 pulse-evoked components based on their respective latency (first and second component) rather than their polarity (negative or positive deflecting components). 78% of electrodes displayed a negative deflection in the first component. The latency at which C1 and C2 Peaks occurred was consistent across electrode contacts, occurring at 21.2 ± 10.1 ms on average for C1 (average ± standard deviation), and 54.2 ± 15.9 ms for C2. The stability of this temporal window supports the importance of relative latency, rather than polarity, for defining the neural components recorded with penetrating electrodes. We also note that our detection was purposefully detecting C1 and C2 in opposing polarities, unlike N1 and N2 components in CCEPs (both negative components). Our modeling approach employs the C1 Peak Amplitude, regardless of its polarity, as the feature of interest best capturing the neural response to stimulation (Figure 2c). Given its fast latency. this component is likely representing the first volley of stimulation-related activity, with minimal contamination by secondary network-wide interactions.

### 4.2. Correlation of derived physiologic and statistical PEP features

Since temporally-defined response components have been previously described as inconsistent in sEEG SPES, we aimed to evaluate a battery of PEP features and their pairwise correlations to examine their stability. Pairwise correlations between all features (C1 and C2 Peak Amplitude, Area, Peak-to-Peak difference, Max Amplitude and RMS-Shape; total of 21 pairs) confirmed very strong correlations among the extracted metrics (correlation coefficients ranging from 0.96 to 0.67, all p-values < 0.0001). The variables more reliably correlating with all other pairs were the C1 Peak Amplitude and the Max Amplitude (correlation matrix marginal mean, excluding diagonal was r = 0.9 for both), while the RMS-Shape displayed the lower correlations with the other variables (marginal mean r = 0.73). The consistency of the correlations validates the selection of the C1 Peak Amplitude feature as the measure of interest for our modeling approach (Figure 4a). To further corroborate this finding, we tested its generalizability to different datasets (Experiments 2 and 3), with different electrodes used as stimulation seeds. The overall agreement between the metrics was preserved, with all pairwise correlations p < 0.0001 also in Experiment 2 and 3. We further validated that SPES targeted to different brain regions led to variable electrical potentials at a given electrode contact (Figure 4b) and showed specific patterns of C1 values across electrode locations. For example, an electrode contact located on the same probe and proximal to the stimulation seed contact, (or an electrode displaying effective connectivity with respect to the stimulation seed location) exhibits an elevated potential during SPES at that stimulation seed. However, that same electrode may display a decreased potential during stimulation at a different stimulation seed. All further analyses and results are based on data from Experiment 1 unless otherwise stated.

**Figure 4.**
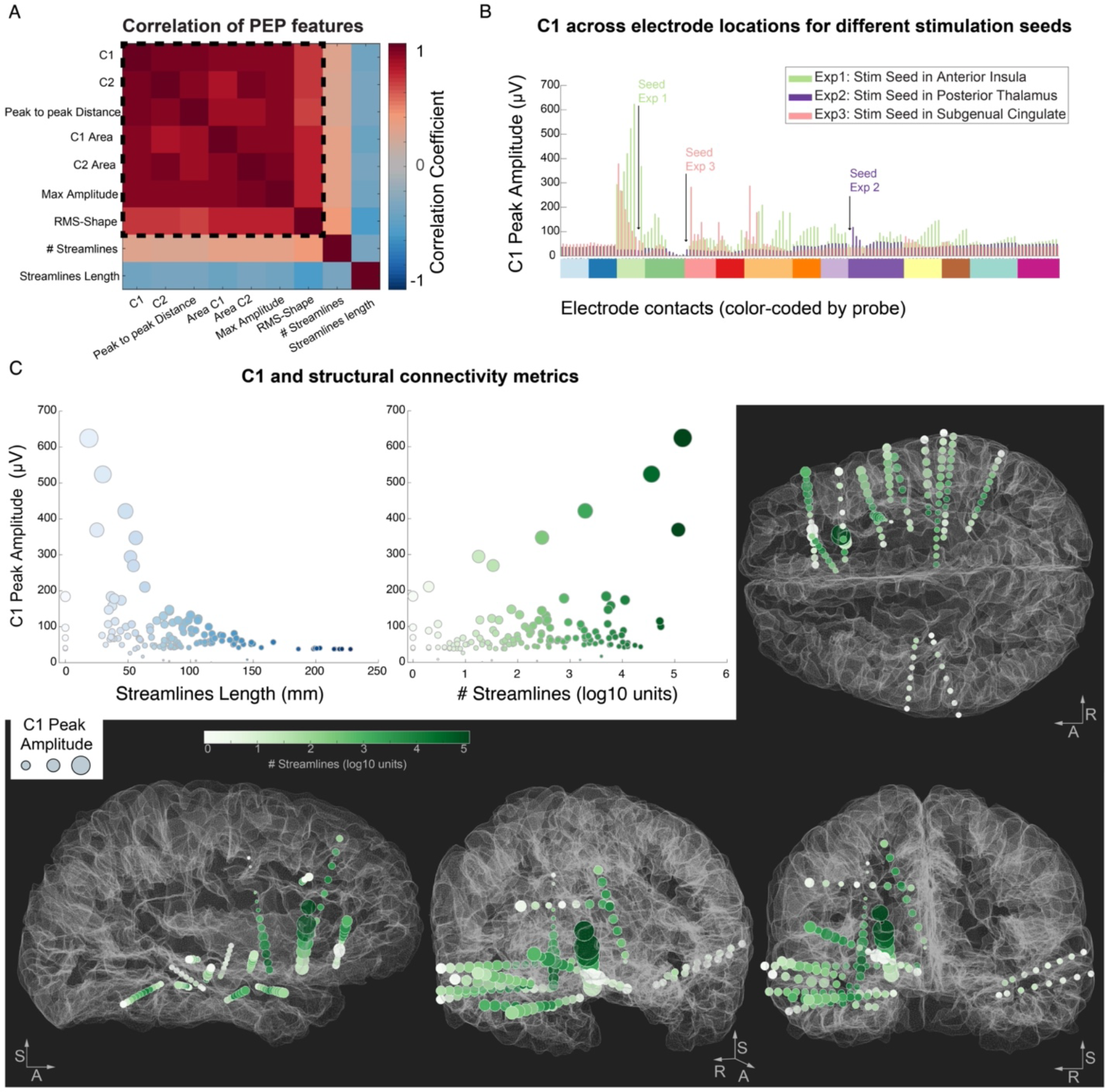
Correlation within PEP features and their relationship with structural information and anatomy. **A**. Correlation matrix for all PEP feature pairs (within the dashed square) and for structural metrics. **B**. Comparison of C1 Peak Amplitude values across the experiments, highlighting how C1 values from the same electrode locations (x axis, color-coded by probe) vary depending on the stimulation seed (location of each seed with respect to the other electrodes is depicted with a black arrow). Experiment 1 and 3 resulted in large C1 values at specific electrode locations (extending beyond the nearby electrodes). Experiment 2 did not elicit large or variable C1 responses beyond the electrodes nearby the stimulation seed. **C**. Correlation of C1 with streamlines metrics in Experiment 1. Left: Streamline Length (average white matter path-based distance between the stimulation seed and each electrode), showing an exponential decay of C1 values with increasing streamline length. Middle: Streamline count (in log10 scale) shows an overall positive relationship with C1 values. Right and bottom: the C1 Peak Amplitude is represented at each electrode location by inflating the size of the sphere by the C1 amplitude (i.e. larger spheres represent larger C1 amplitudes). The electrode locations are color-coded by the Streamline count between each electrode location and the stimulation seed to display the presence of large number of streamlines passing through the electrode locations in darker green colors. The R-A-S axis (right, anterior, superior) are included for reference.

### 4.3. Concurrence among structural and effective connectivity modes

Cross-modality studies have demonstrated moderate similarity in connectivity measured by tractography with the occurrence and magnitude of PEPs (Adkinson et al., 2022; Crocker et al., 2021). However, the nature of the relationship between tractography and SPES-based connectivity remains incompletely understood. Thus, we examine, in the context of sEEG SPES, how connectivity strength and path length correlate with PEP features. Pearson correlations were conducted in a pairwise manner between PEP metrics and tractography. The connectivity likelihood was computed by counting the total number of streamlines for all stimulation seed and target electrode contact pairs (and applying a base 10 logarithmic transformation to correct for skewness). This metric (# Streamlines) demonstrated a positive correlation with all PEP features (Figure 4a). The correlation values ranged between 0.25 and 0.40, thus considerably lower than the correlation obtained within the PEP features, but nonetheless statistically significant (all p-values < 0.01). The second structural metric examined was the average length (in mm) of the streamlines connecting each electrode location to the stimulation seed (Streamlines Length), capturing the distance from the stimulation seed based on the white-matter pathways connecting them. All PEP features demonstrated a moderate negative correlation with the average Streamline Length (Figure 4a), with correlation coefficient values ranging between -0.58 and -0.33 (all p-values < 0.0001). Thus, electrodes farther away from the stimulation seed display smaller PEP response features. Interestingly, the most negative correlation (r = -0.58) was found between the average streamline path length and the RMS-Shape, suggesting that PEP waveforms measured at farther locations from the stimulation seed display less consistent PEP waveform across SPES repetitions.

Finally, we evaluated the C1 peak amplitude feature and its relationship to the tractography-estimated metrics. The C1 component was explained by an exponential decay in amplitude with increasing average streamline length (Figure 4c left), with a half-life of 44.1 mm, obtained by fitting an exponential decay function of the form: *y* = *ae*^−*l*/*b*^ to the streamline length (represented by *l*) and the C1 Peak Amplitude values (*y*). On the other hand, we observed a positive but weak correlation between C1 and streamline count (r = 0.29, p<0.001), with C1 amplitudes above ∼150 µV displaying a very consistent association (Figure 4c, middle). Thus, while the likelihood of structural connectivity relates to the PEP response magnitude, the relatively low correlation values are indicative that the variation in C1 amplitude across brain locations cannot be fully captured by this information alone (Figure 4c, right and bottom).

### 4.4. 3-dimensional topology optimization model resolves conductivity map across electrode contact locations

The electric potentials of the first temporally-defined PEP component, C1, detected at each electrode contact during SPES at the stimulation seed used in Experiment 1 (Figure 1c) were used as input data to the optimization model. Across model optimization iterations, a conductivity map explaining known voltage behavior with minimum error is constructed, representing a general nodal connectivity path. The proposed model achieves an overall average error of 14% between measured and simulated potentials across all electrode locations (median 6%, ranging from 0 to 123%; Figure 5a). Notably, the model does not use or require anatomical constraints. When considering the electrode locations with the largest error, three electrode locations stood out as outliers (above 100%). Of these, two were located on the same probe as the reference electrode and had small C1 values (3 and 8 µV) that the model could not properly account for, leading to negative potential estimates in those locations. The overall worst electrode (123% error) was located in the right inferior frontal cortex, spanning *pars triangularis* and *orbitalis* and less than 1 mm from the gray-white matter boundary. Overall, inspection of the electrodes displaying the larger errors revealed that the model performed less accurately on locations where the C1 Peak Amplitudes varied considerably between neighboring electrodes (Figure 5a).

**Figure 5.**
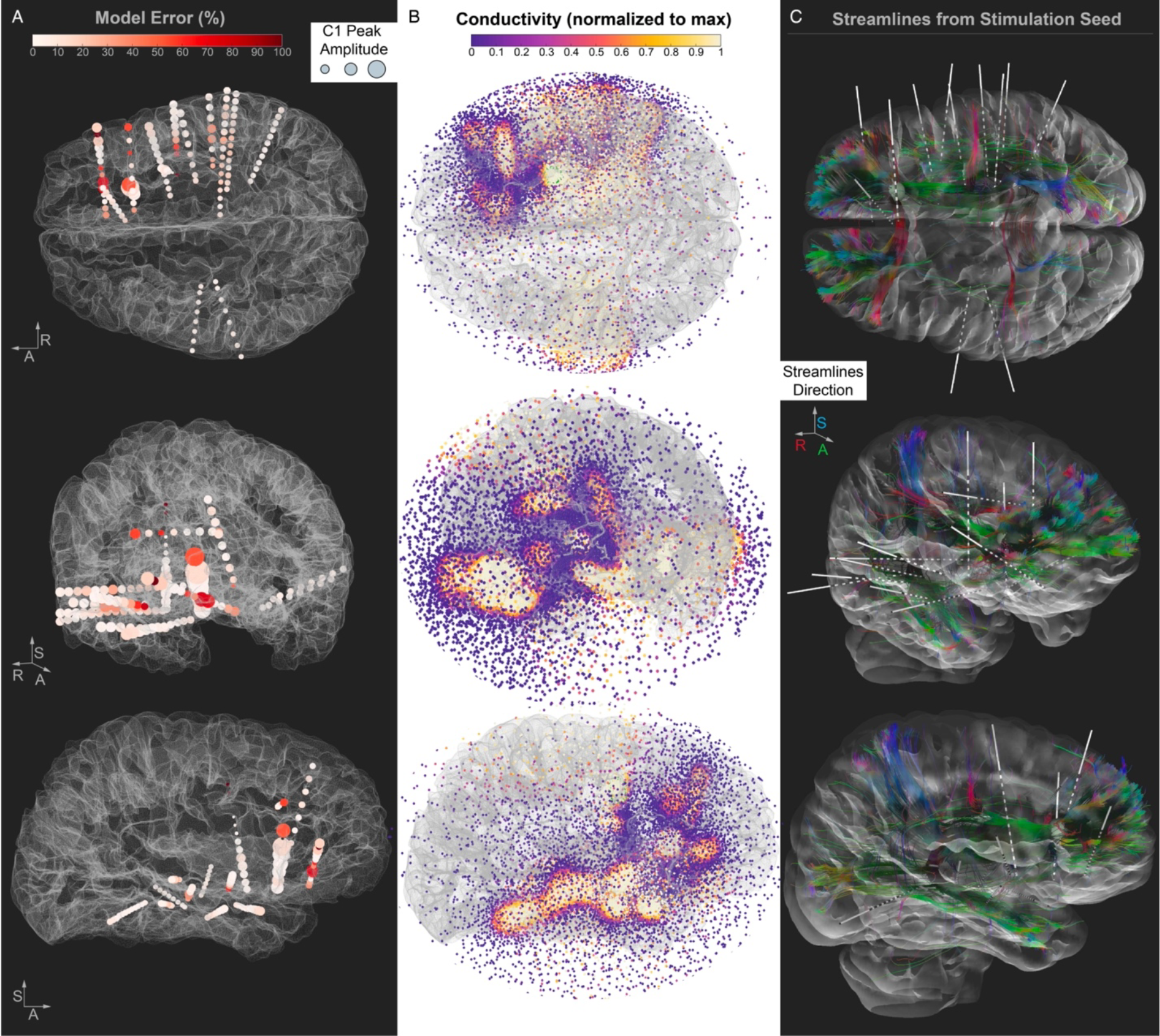
Visualization of model results and structural information. **A**. The Model Error (% difference between C1 value and modeled potential) is plotted at each electrode location and color coded by a white to red scale. The C1 Peak Amplitude is represented at each electrode location by inflating the size of the sphere by the C1 amplitude (i.e., larger spheres represent larger C1 amplitudes). The R-A-S axis (right, anterior, superior) are shown as a reference. **B**. The conductivity map solution obtained by the model is represented by spheres at the model mesh locations, color coded by conductivity. The conductivity map solution achieved by the model is overlaid on the brain surface (same orientations as A) to facilitate a visual comparison of the conductivity map and the other panels. **C**. Streamlines between the stimulation seed and all electrode locations, color coded by the direction of the streamlines. The stimulation seed electrode is represented by an inflated green sphere. Figure generated using DSI Studio (https://dsi-studio.labsolver.org/) where deterministic fiber tracking (Yeh et al., 2013) was conducted using modeled diffusion data from our FSL processing pipeline.

As depicted in Figure 5b, the conductivity values across the model domain (i.e., the sphere used to model the space around the electrodes) vary, supporting propagation of conductivity in spatially preferred directions reaching electrode probes. We investigated the different model parameters, and we noted that the absolute value of the source potential does not significantly affect model accuracy so long as the value is larger than the maximum C1 value. A constant potential *V*_0_ = 700 μV was placed on the boundary of the source electrode, a source which exceeds the maximum C1 value of 624.57 μV. As expected, increasing the source potential further leads to larger void regions (low θ_p_ and thus conductivity values) immediately around the source electrode. The model is unconstrained by anatomy and has many degrees of freedom in addition to the free solver parameters discussed in Section 3. We found that a critical constraint was accounting for the reference potential *V*_*ref*_ in the objective function. A preliminary model that did not track the reference potential had very poor accuracy, with a 43% mean error (median 7%, ranging from ∼0 to 1362%). The four electrodes with the largest error were located on the reference probe and were among the smallest in C1 value across the dataset. Explicitly probing the reference potential in the model at the location of the experimental reference improved the maximum error by an order of magnitude (1362% to 123%).

Next, we evaluated the similarity between the results of the model and the structural connectivity information offered by diffusion-based tractography. Despite the lack of any information on underlying anatomy, we observe a visually identifiable relationship between the 3D conductivity map and the tractography streamline map (Figure 5b, c). We interpolated the conductivity values and rendered them into a volume, allowing for a more careful overlap between the model results and the white matter pathways in a specific sagittal plane encompassing prefrontal and temporal cortex (Figure 6a). An obvious caveat to the interpolated conductivity values is that locations farther away from the sampled cortical locations might diverge from the structural information. For example, parietal, posterior cingulate, and occipital connections cannot be captured in our results due to the lack of electrodes sampling from these regions in our dataset, as discussed above. Overall, the model would likely resemble the structural information more closely if applied to a dataset with a more distributed cortical sampling, offering a better triangulation of the voltage values.

**Figure 6.**
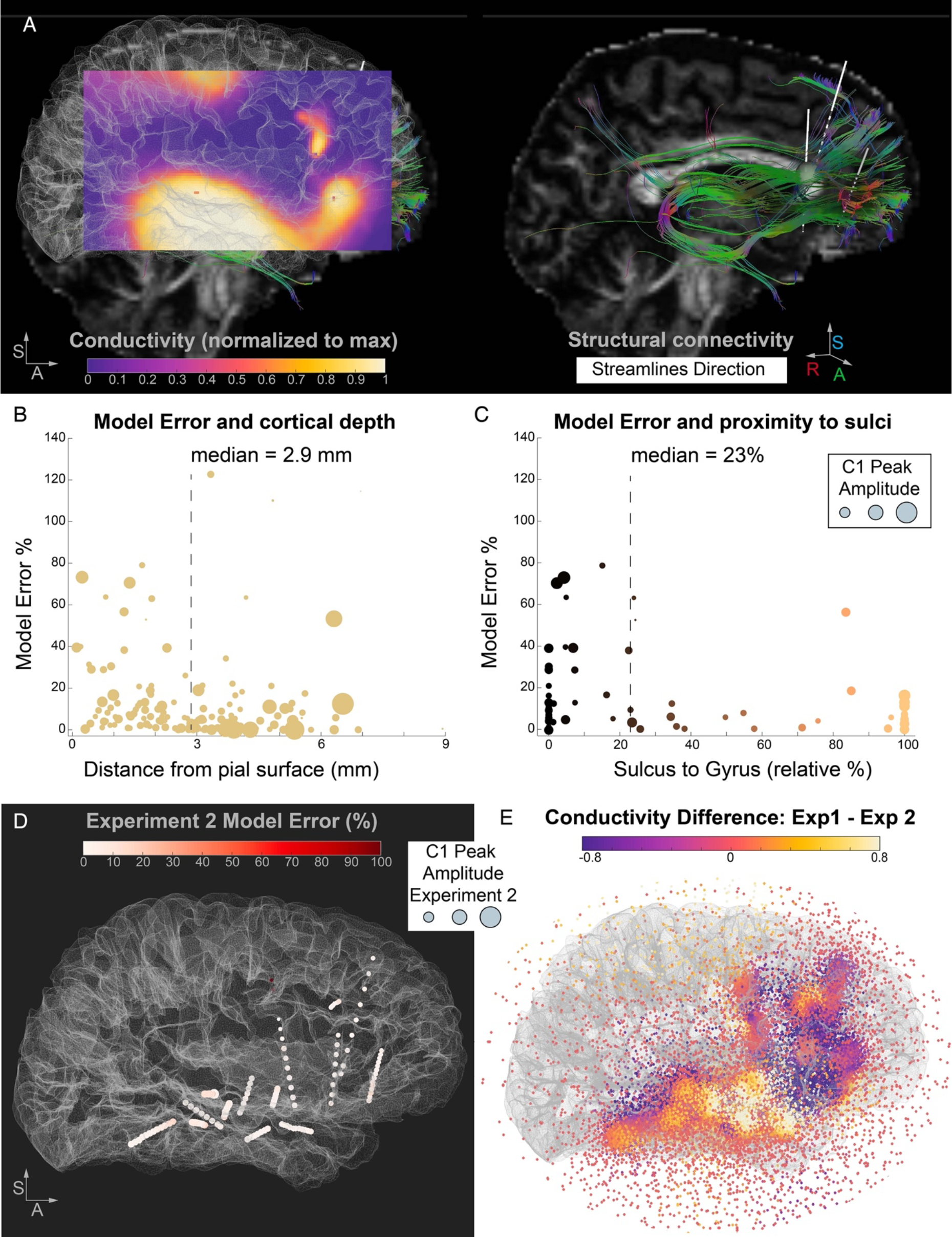
Model error dependency on anatomical features and conductivity map differences between different stimulation seeds. **A**. The conductivity map solution obtained by the model is interpolated and represented by a sagittal slice color coded by conductivity. The map solution is overlaid on the brain surface (same orientations as panel B) to facilitate the visual comparison of the conductivity map and the streamlines. On the right, the streamlines between the stimulation seed and prefrontal electrode locations are shown, color coded by the direction of the streamlines. The stimulation seed electrode is represented by an inflated green sphere. **B**. Relationship between model error and distance from cortical surface. The model error is significantly larger at electrode locations closer to the pial surface versus farther away (closer to the white-gray boundary). Dashed line depicts the median distance value used to perform the split of the electrodes into subgroups. **C**. Relationship between model error and electrodes in gray matter (subset of 69 electrodes). Electrodes close to a sulcus (versus a gyrus, relative proportion) have higher model errors, indicative of the effect of folding patterns on the presence of large voltage gradients. **D**. Experiment 2 Model Error (% difference between input C1 value and modeled potential) is plotted at each electrode location and color coded by a white to red scale. The C1 Peak Amplitude is represented at each electrode location by inflating the size of the sphere by the amplitude values (i.e. larger spheres represent larger C1). The stimulation seed in Experiment 2 (posterior thalamus) did not elicit large or variable C1 Peaks across the electrodes and the model for Experiment 2 achieves lower errors with respect to Experiment 1. Panel E. Comparison between the conductivity solutions for the two models (Experiment 1 versus Experiment 2), highlighting spatial locations of disparate effective connectivity as a function of stimulation seed.

While most data are modeled accurately, some electrode locations displayed significantly large error values, and we next focused more in depth on factors that could explain the observed variability in the error rates. As discussed above, one factor contributing to the error distribution across brain locations is the presence of drastic differences in the potentials of neighboring electrodes (Figure 5a, large C1 Peak Amplitude differences among electrodes on the same probe), lowering the numerical model performance. Large potential differences across isolated small regions of space (i.e., the neighboring electrodes) imply large gradients that need to be associated with localized small conductivity values. Therefore, large voltage gradients translate into potentially large conductivity spatial gradients that are computationally demanding to address during the optimization process. The model, being naive to the underlying anatomy, may be unable to fit a spatial solution that fully accounts for sharp condition changes. However, from an anatomical perspective the complex cytoarchitecture of the human brain may contribute to observations of large differences in recorded potentials from neighboring electrode contacts. Specifically, sulcal landmarks have been recently described as natural barriers to electrical activity propagation (Maharathi et al., 2019) and stimulation-induced electrical fields modeled variably based on the positioning of a stimulated electrode relative to sulcal orientation (Alonso et al., 2023).

### 4.5. Location of high model error relates to proximity to complex anatomy

To test whether the electrode locations displaying high error can be tied to anatomical organization, we calculated and compared various anatomical features for each electrode location, including the distance to cortical surface, the distance to gray-white matter boundary, and the likelihood of an electrode contact being close to a sulcus versus a gyrus. We tested the association between these anatomical features and model error by performing median splits of the dataset based on each anatomical feature. The distance from the cortical surface significantly modulated the error, with electrodes closer to the pial surface showing higher model errors (8.2% vs. 3.7%, z = 3.49, Wilcoxon Rank Sum Test, p < 0.001, Figure 6b). Finally, we focused on the electrodes localized in gray matter (i.e., electrodes with coordinates localized between the pial surface and the gray-white boundary, n = 69 out of 143) and asked whether an electrode’s proximity to a sulcus affects model performance. We found that electrodes closer to a sulcus demonstrate larger model error (17% vs. 10%, z = 2.9, p = 0.004, Figure 6c). This result suggests that anatomical features can explain some variability in the model error, supporting the idea that complex cortical folding patterns (which would impact dipole geometry) in the proximity of the electrodes might be responsible for sharp transitions in PEP features that are not easily accounted for by a passive modeling approach.

Because tractography algorithms are known to perform poorly in tracking sulcal regions (Chen et al., 2013; Reveley et al., 2015), we investigated the relationship between structural connectivity strength and model error. As expected, electrode contacts displaying high error also demonstrated weak to no structural connectivity. Beyond sulcal banks, tractography also poorly tracks gray matter regions, due to lower diffusion anisotropy in and around neuronal cell bodies and axon terminals. To further parse the relationship between model error and tractography, we adjusted our ROI masks for gray matter regions. Electrode contacts identified as residing in gray matter were rerun for each stimulation seed using a larger radius spherical mask (3 mm) to aid the tractography algorithm in detecting nearby streamline terminations, as diffusion in gray matter shows much weaker directional consistency than in white matter. We then compared the difference between connectivity strengths at the original and new ROI spherical mask sizes with the model error. We found that electrode contacts located in gray matter regions with increased connectivity likelihood (n = 29) did not have a significantly different model error with respect to electrode contacts with no improvement in streamline count (n = 40; z = 0.6261, p = 0.53). Taken together, these results suggest that our model can resolve conductivity to gray matter regions, unlike tractography, while displaying a lower accuracy for paths near or in sulci.

### 4.6. Generalizability to a second dataset

The above modeling results were obtained from data in Experiment 1. As a final step, we applied the same approach to the dataset from Experiment 2, where the stimulation seed was in the posterior thalamus. Overall, the C1 Peak Amplitudes in this dataset were smaller and less variable across electrode locations (Figure 4b, purple bars). From an electrophysiological perspective, this can be ascribed to the stimulation seed not displaying a strong and specific connectivity pattern to any of our sampled locations. The model error for Experiment 2 was considerably lower than Experiment 1 (average error 1.7%, with a median of 5.5% and ranging from ∼0% to 126%; Figure 6d). Thus, the model found a solution able to more accurately represent this homogenous dataset (i.e., less variable C1 Peak Amplitude values across brain locations). Interestingly, the top two worst errors in this model (110 and 126% error) were the two electrodes closest to the reference location, associated with very small C1 Peak Amplitudes (4 and 6 μV), mirroring the result obtained in Experiment 1. This generalizes the previous finding that the conductivity topological information in proximity of the reference electrode is less likely to be appropriately modeled and highlights the need of careful consideration when selecting the reference electrode for intracranial recordings. Overall, the results obtained validate the interpretation that our approach can resolve different conductivity maps and that the error can be used as a heuristic to identify locations associated with large gradients, as in cases where the stimulation seed shows a high effective connectivity with a specific subset of locations. This indeed leads to a higher variability in the C1 Peak Amplitudes, as found in Experiment 1, but not in Experiment 2. The direct comparison of the two conductivity maps (Figure 6e) highlights the prefrontal and temporal regions displaying the highest effective connectivity differences.

## 5. Discussion

In this paper, we introduce a 3-dimensional conductivity model capable of predicting brain connectivity among functional networks. Provided with electric potentials recorded in-vivo from sEEG electrodes, our optimization model resolves a conductivity solution via passive propagation at nodes spanning spatially disparate distances to articulate pathways of brain connectivity. The proposed model remains naïve to anatomical organization, such as sulcal barriers. The electric potentials guiding our model are reliable and early pulse-evoked potential response components to single pulse electrical stimulation in sEEG, termed here C1. We evaluate the correlation of derived components C1 and C2 with other established PEP features and further explore the structural and effective connectivity relationship from high-resolution tractography data. Noting these relationships, we identify and prospectively associate model error with anatomical features, providing insight to model performance and interpretation.

Understanding the behavior of electrical activity in the brain remains an ongoing goal in both clinical and research settings. Computational modeling approaches have consistently advanced toward describing conductivity in relation to induced stimulation fields and identifying conductivity of specific neural tissue structures. However, using conductivity as a lens for modeling functional brain connectivity remains less explored. Alternative methods for constructing conductivity maps in the human brain, such as electrical impedance tomography, are limited by confounds such as skull impedance (Oh et al., 2009). Other techniques, including magnetic resonance electric properties tomography, extract conductivity at high frequencies from magnetic resonance signal read-outs, though these approaches are dependent on scanner frequency and signal phase processing (Borsic et al., 2016; Voigt et al., 2011). Complicating estimations of conductivity in the brain is the fact that conductivity in neural tissue varies with tissue type (gray matter, white matter, CSF) and composition (McCann et al., 2019). A prominent component across neural tissue types is water, which may be associated with electrical properties in biological tissue based on concentration (Schepps and Foster, 1980). This relationship served as a basis for conductivity tensor imaging, combining water maps with diffusion MRI data (Marino et al., 2021). Critically, these methods rely on the strong assumption that water concentration is determinant of conductivity. The relationship of water molecules and brain conductivity is especially interesting in light of our model. Our initial inclusion of diffusion-based tractography data was to explore the association between structural and effective connectivity metrics. However, we additionally observed a visually identifiable relationship between the volumetric topology conductivity map and the tractography streamline map. These similarities exist despite a lack of input information on water behavior to our model. Such results optimistically support a relationship between water concentration and conductivity in neural tissue elements, and future work should continue to explore this relationship more intimately.

The observed similarity between streamline and conductivity maps motivated subsequent investigation into possible anatomical explanations for model error. While most electrode contacts are resolved with minimal error, a subset of contacts present high error percentages. Extending beyond physics-based reasoning for higher error (see Methods section 4.4), the existence of structural barriers in the brain may play corresponding roles. We focus on the existence and proximity of sulcal bounds, since these areas present well-documented difficulties for tractography-based connectivity methods (Van Essen et al., 2014). Specifically, tracking algorithms bias toward termination points along gyral crowns, failing to quantitatively account for white matter fibers crossing the white-gray matter boundary (Schilling et al., 2017). As anticipated, our results support that sulcal bounds may represent one anatomical predictor for model error. Fiber connectivity densities have been shown to be elevated in gyral crowns compared to sulcal barriers, and fiber bending over the distance of even 1 mm can occur almost orthogonally along these proximal walls (Schilling et al., 2017). Thus, convoluted organization of the cerebral cortex into sulci and gyri and the accompanying variation in neuron orientation introduce anatomical complexities that may underlie some degree of model error. Additionally, we observe high model error at electrode contacts located on the same probe as the reference electrode. One important consideration in brain stimulation experiments is selecting the location of an intracerebral reference electrode. Typically, electrodes in well-insulated, deep white matter locations are preferred, though regardless of location, gleaning meaningful information from surrounding electrodes proves difficult. In our model, the high error observed at contact locations proximal to the reference demonstrates an inability to accurately model these locations. While selecting the optimal reference location, investigators should keep in mind the consequent loss of information in nearby regions to best avoid designating an anatomically intriguing region as the reference.

The proposed method models passive propagation of electrical signals. In biological tissue, both active and passive signal transmission occurs, where the latter sums electric potentials along the temporal domain with spatial-dependency (Elmslie, 2021). While passive electrical propagation may describe synaptic and receptor potentials, actively propagated signals arise from action potentials (Purves et al., 2001). Given that our model focuses on passive propagation, electrode contacts displaying high error may indicate a dominant role played by active potential summation effects. We can then assume that at specific nodes, higher conductivities are not accounted for by passive principles. In Experiment 2, using a stimulation seed in the thalamus, C1 Peak Amplitudes displayed little variation across stimulation sites. In pairing, our conductivity map was resolved with minimal error, supporting that electric potentials may have resulted from volume conduction effects. This is in comparison with higher error observed in Experiment 1, where stimulation of the anterior insula resulted in regionally dependent C1 amplitudes, and our conductivity map presented higher error. Thus, where C1 amplitudes were selective in response size across regions in the insula, the thalamic stimulation seed behaved in a non-selective manner, nearly preserving C1 response independent of spatial location. From this, we proffer that our model is informative about hyper-selective paths.

A secondary objective of the present study was to investigate the derived PEP features with tractography-based metrics, probing the relationship between functional effective and structural connectivity. We first evaluated statistical and physiologic PEP measures against one another and observed significantly strong correlations across all variables, validating our C1 and C2 detection algorithm. Then, we compared two tractography-based metrics, number and length of streamlines, against each PEP feature. For the number of streamlines, we reported stable yet weak correlations with PEP features, with the strongest positive association to RMS-Shape. This result supports that streamline count correlates with consistent PEP responses across stimulation repetitions, suggesting a stronger structural relationship between seed and target electrode location may tie with stable effective responses. Separately, we observed moderate negative relationships between path length and all PEP features, with the strongest effect also from PEP shape consistency. As distance between stimulation seed and target electrode contact increases, weaker consistency in waveform shape is preserved. Such influence of distance on SPES-induced evoked responses has been previously documented, diminishing the observed relationship between stimulation response and diffusion-based tractography connectivity measures (Crocker et al., 2021). For the C1 component amplitude, reliable but weak correlations were observed with both streamline count and path length. While an exponential decay function describes decreasing C1 peak amplitude with increased distance, the relationship between C1 and number of streamlines appeared less obvious. Overall, our findings are consistent with previous work and support that while structural connectivity measures can stably track with PEP features, effective connectivity, especially C1 amplitude, cannot be fully captured from tractography-predicted connectivity. One possibility is that, given the limitations of structural connectivity as described above, C1 Peak Amplitude may more closely match underlying anatomical connectivity. This will need to be studied using tract-tracing in nonhuman animal models. Future studies are needed to continue expanding on the inter-relations across connectivity types.

Several limitations of the current work are notable. As previously mentioned, the proposed model solely captures passive electrical signal propagation. Despite the informative nature of error percentages, without considering both active and passive propagation, the proposed model cannot account for the physicochemical mechanisms underlying excitability and inhibitory tissue activations. Additionally, the input parameters used in model optimization are not identical to those used for the SPES waveform. Since modulation of stimulation parameters influences evoked potential response morphology (Paulk et al., 2022), the generated conductivity map may only partially represent conductivity propagation from the stimulated seed locations. For simplicity, we implemented a spherical volume mesh rather than a mesh topology based on brain geometry. Future work will employ anatomical boundary constraints. As patient data was collected from sEEG electrodes, spatial sampling was limited to the surgical implant strategy. In the present paper, electrode coverage primarily targeted the right hemisphere, resulting in sparse sampling of the left hemisphere. As such, our model is limited in its ability to resolve conductivity propagations into the contralateral hemisphere. Future studies will further exploit experimental data by considering the latency of the recorded signals at the electrodes with respect to SPES. While the introduction of the time coordinate may increase the computational complexity, it will also provide additional constrains to the optimization procedure, potentially increasing the accuracy of the resulting conductivity maps. Lastly, input intracranial recording data was collected from a single patient. Thus, this model was constructed on individual patient data and has not been tested on other subjects. To promote further generalizability and increased spatial sampling, more patients with additional electrode coverage should be evaluated. Nonetheless, the topology optimization model presented provides a necessary introduction into using intracranial recording data for reconstructing conductivity paths in the human brain to advance our understanding of functional brain networks.

## Acknowledgments

We thank the patient that took part in the experiments. We thank Stephen K. Sanders for his assistance with COMSOL. This work was supported by the NIH (R01-MH127006 to K.R.B. and R21-MH127842 to A.J.W.) and by the McNair Foundation (S.A.S.). This material is based upon work supported by the National Science Foundation Graduate Research Fellowship under Grant No. 1842494 (to W.S.). Any opinions, findings, conclusions, or recommendations expressed in this material are those of the authors and do not necessarily reflect the views of the National Science Foundation.

## Author contributions

Conceptualization: E.B. A.A. K.R.B. S.R.H. J.A.; Investigation: I.A.D. J.A. E.B. A.W.; Formal analysis: W.S. A.A. M.C.E. I.A.D. J.A. E.B.; Visualization: W.S. I.A.D. M.C.E. J.A. L.M. E.B.; Writing: I.A.D. W.S. M.C.E. E.B. (original draft) and A.A. K.R.B. S.R.H. A.W. G.P.B. S.A.S. (review editing).

## Appendix A

**Figure A1.**
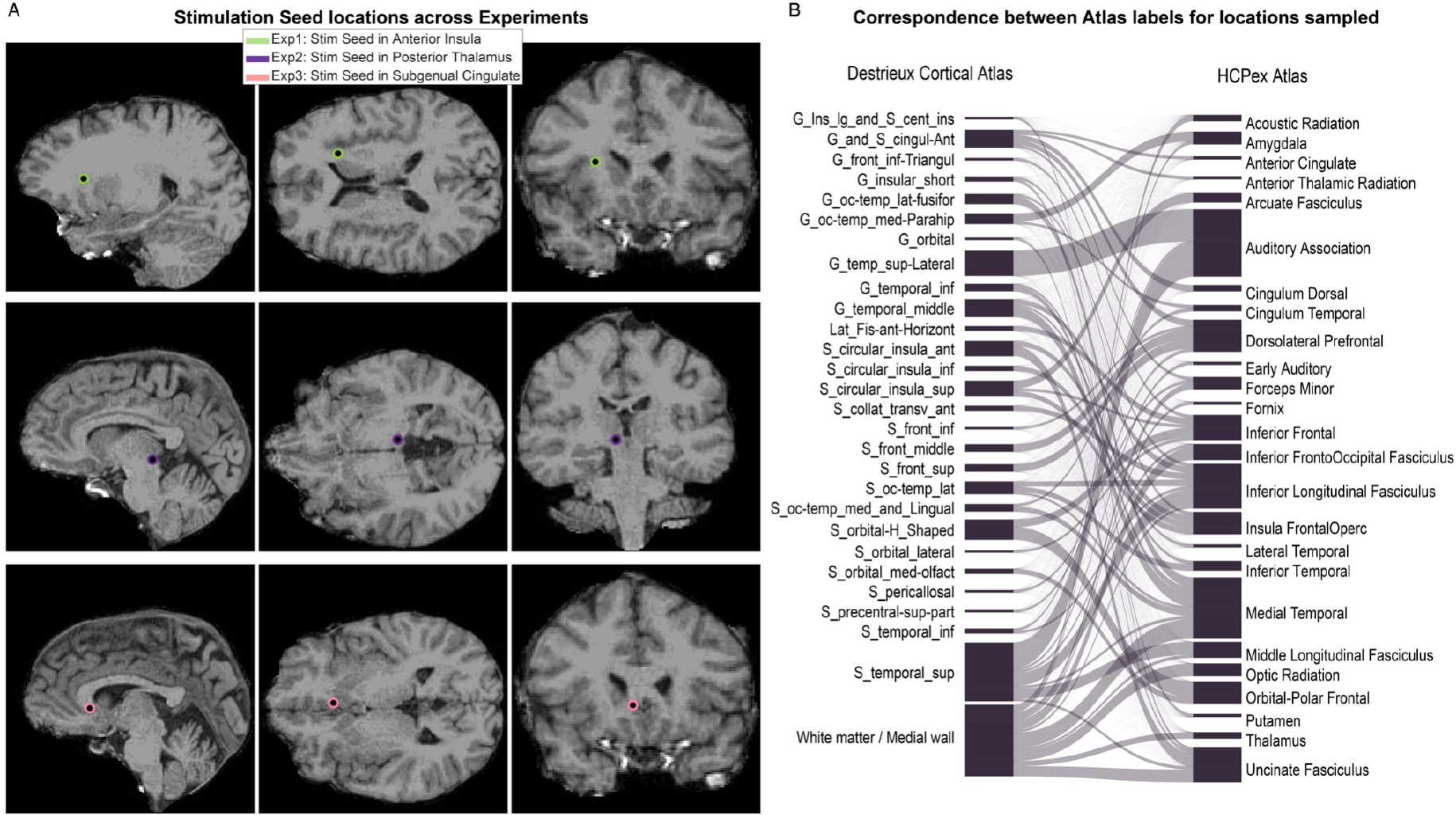
Electrode locations used for stimulation (stimulation seed) and cortical parcellation estimates of all electrodes. **A**. sagittal, axial and coronal view of the location of each stimulation seed (Experiment 1: white matter adjacent to anterior insula; Experiment 2: posterior thalamus, Experiment 3: subgenual cingulate cortex). **B**. Bipartite plot mapping the anatomical parcellation estimates obtained for each electrode across the two atlases employed in the analysis (Destrieux Cortical Atlas and modified HCPex Atlas).

